# Asymmetric survival in single-cell lineages of cyanobacteria in response to photodamage

**DOI:** 10.1101/2022.04.14.488368

**Authors:** Jian Wei Tay, Jeffrey C. Cameron

## Abstract

Oxygenic photosynthesis is driven by the coupled action of the light-dependent pigment protein complexes, photosystem I and II, located within the internal thylakoid membrane system. However, photosystem II is known to be prone to photooxidative damage. Thus, photosynthetic organisms have evolved a repair cycle to continuously replace the damaged proteins in photosystem II. However, it has remained difficult to deconvolute the damage and repair processes using traditional ensemble approaches. Here we demonstrate an automated approach using time-lapse fluorescence microscopy and computational image analysis to study the dynamics and effects of photodamage in single cells at sub-cellular resolution in cyanobacteria. By growing cells in a two-dimensional layer, we avoid shading effects, thereby generating uniform and reproducible growth conditions. Using this platform, we analyzed the growth and physiology of multiple strains simultaneously under defined photoinhibitory conditions stimulated by UV-A light. Our results reveal an asymmetric cellular response to photodamage between sibling cells and the generation of an elusive subcellular structure, here named a ‘photoendosome’, derived from the thylakoid which could indicate the presence of a previously unknown photoprotective mechanism. We anticipate these results to be a starting point for further studies to better understand photodamage and repair at the single-cell level.

## Article

Photosystem II is a pigment-protein complex that resides on the thylakoid membrane and catalyzes the light-dependent oxidation of water and reduction of plastoquinone. The heterodimeric core of PSII is comprised of the D1 and D2 proteins which bind essential redox-active cofactors^1^. However, the D1 and D2 proteins are prone to oxidative modification by reactive oxygen species (ROS) that are generated at multiple sites of PSII^2^. Therefore, a repair cycle that entails the partial disassembly of damaged PSII, followed by replacement of the damaged D1 is required to maintain a population of active PSII proteins^3^.

The steady-state level of functional PSII in a cell is determined by the relative rates of synthesis, degradation, dilution through cell-expansion and division, damage, and repair (Figure 1a). Consequently, when cells are exposed to excess light such that the rate of photodamage exceeds the rate of synthesis and repair, photochemical efficiency and biomass production is reduced in a process termed photoinhibition^4^. Early experiments proposed at least two distinct modes of photoinhibition based on the action spectrum of photodamage, with both chloroplasts in plants and cyanobacteria exhibiting higher sensitivity to UV compared to visible light^5,6^. However, there are significant differences in the protein constituents and molecular mechanisms of photosystem assembly, photodamage, and repair process between plants and cyanobacteria that remain to be resolved^3^

**Figure 1.**
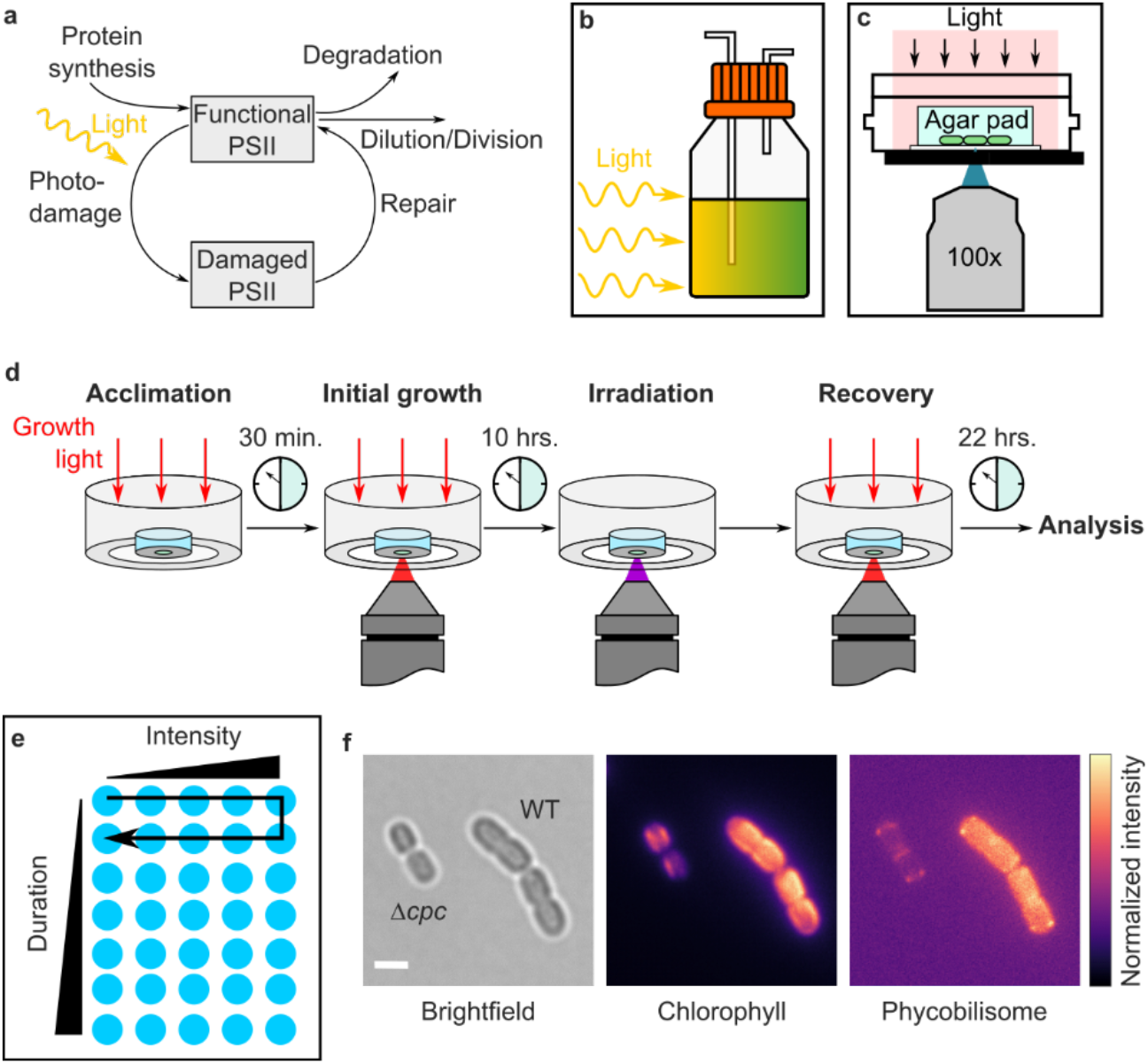
ystem description. **a**, Illustration of the processes determining the concentration of functional PSII in the cell. **b**, In bulk cultures light penetration is attenuated due to cell-cell shading, resulting in uneven light exposure. **c**, In our approach, cells are grown in a single 2D layer on a microscope, allowing uniform and controlled delivery of light for photosynthetic experiments. **d**, Illustration of the different phases of the experiment. Cells were initially incubated for 30 minutes to allow for acclimation to the microscope environment prior to imaging with an inverted 100x objective (1.45 N.A.). They were then filmed for 10 hours to obtain initial growth measurements. The cells were then irradiated with pulses of UV-A (395/25 nm) light, then imaged for a further 22 hours to observe recovery. Growth light (610-650 nm LED; ∼157 μmol photons m^-2^ s^-1^) was provided from above during acclimation, growth, and recovery. **e**, The duration and intensity of the irradiation pulse were varied over 35 physically separated x-y locations on the same agar pad, allowing different experimental conditions to be carried out simultaneously. **f**, Representative images showing the transmitted light (brightfield), the chlorophyll fluorescence, and the phycobilisome fluorescence of WT and Δ*cpc* strains. The color bar indicates the scale of the fluorescence intensities, which were normalized to the same relative scale for each image. Scale bar is 5 μm.

Current approaches used to study photoinhibition in cyanobacteria require growing cells in bulk cultures to provide sufficient cellular material for biochemical and biophysical assays^7–10^. However, the incident light quality and quantity received by individual cells within the culture vessel is greatly impacted by mixing and light penetration due to cell-cell shading^11^. While use of high-intensity light can overcome some of these challenges^12^, changes in light penetration resulting from dynamic alterations in cell density or cellular pigment content can lead to heterogenous populations and physiological states during studies of photoinhibition (Figure 1b). Furthermore, standard methodologies for measuring photoinhibition rely on the use of protein translation inhibitors chloramphenicol and lincomycin to inhibit competing repair processes, despite the potential for these compounds to result in side effects with unintended consequences. For example, chloramphenicol was recently shown to artificially enhance photodamage of PSII, likely through the production of superoxide^13^.

To overcome these limitations, we have developed an automated imaging approach using long-term time-lapse fluorescence microscopy to film cyanobacterial cells under optimal growth and stimulated photoinhibitory conditions. Here, cells are grown in a two-dimensional layer (Figure 1c), thereby eliminating shading effects and allowing cells to be exposed to highly reproducible growth conditions. In our approach, cells are first pre-illuminated with a defined monochromatic light source to enable steady-state growth before exposure to controlled pulses of light. These pulses stimulate photoinhibition, thereby eliminating the need for protein translation inhibitors. Subsequently, cells are returned to growth light to enable recovery from photoinhibition. Multi-channel transmission and fluorescence images are acquired over the time course to provide quantitative information on cellular morphology and ultrastructure, as well as native pigment-protein complex and thylakoid membrane dynamics. Custom computational image processing and cell-tracking tools allow the growth rates of individual cells to be tracked over multiple generations with sub-cellular resolution^14^. Using this integrated imaging-computation system, we simultaneously analyzed the growth and physiology of multiple strains co-plated on the same substrate while titrating photoinhibitory conditions. Therefore, our system enables both experimental and control strains to be directly compared under identical conditions and provides a robust new methodology for future photosynthesis research.

## Results

### Experimental description

Here we report on a proof-of-principle experiment using our new platform to induce photodamage in actively growing cyanobacterial cells. In this work, we compared the response to photoinhibitory conditions in a wild-type (WT) fast-growing and high-light tolerant marine cyanobacterium, *Synechoccocus sp. PCC 7002* (hereafter PCC 7002)^15^, and a mutant strain lacking the *cpcB-F* operon (hereafter Δ*cpc*), which encodes for part of the phycobilisome rods^16^ (Figure 1f). When coupled to PSII, phycobilisomes act as an energy funnel, directing light energy absorbed by the rods to the central core of the phycobilisome and the PSII reaction center^17^. Reduced absorption cross section of PSII resulting from antenna truncation leads to light limitation and reduced growth compared to the WT under typical growth conditions^18–20^, making it difficult to directly compare the physiology and light response of these strains in bulk culture. Recently, adaptive laboratory evolution studies under high-light conditions selected for strains with reduced levels of antenna^21^. These studies provided the motivation to directly compare growth and photoinhibition of WT and Δ*cpc* strains in 2D cultures.

An outline of the experiment is shown schematically in Figure 1d. The two strains were grown overnight in separate liquid cultures, then mixed and spotted onto an agarose pad (0.5% w/v A+ media) (Extended Data Figure 1). The Δ*cpc* strain was previously shown to be mechanically sensitive to agarose concentrations >0.5%^18^. The pad was placed into a glass-bottomed imaging dish (μ-Dish, Ibidi), which was then inserted into a temperature-controlled chamber on a mechanized fluorescence microscope (Nikon TiE) and allowed to incubate for 30 minutes to acclimatize the cells to the new environment. During this incubation period, the cells were illuminated with a red LED (610-650 nm) at an intensity of ∼157 μmol photons m^-2^ s^-1^ (Lida Light Engine, Lumencore), which was previously determined to be optimal for microscopic growth of PCC 7002^18,22^. Following this initial incubation period, the cells were imaged for 10 hours. This 10-hour period was chosen to allow both strains to undergo at least one cell division, thereby ensuring they were out of lag phase. Following this growth phase, the cells were exposed to 395 nm light (UV-A), which was recently shown to directly cause light-induced damage of the D1 protein in plants^23^. To test multiple conditions simultaneously, we programmed the microscope to irradiate cells in 35 physically separated x-y locations on the agarose pad with pulses of the UV-A light of varying duration between 0.5 - 3.5s, and intensity (Fig. 1e). The intensity was further controlled by programmatically changing the laser power settings between 5 - 85%, corresponding to ∼1.31 - 19.7 mW over an area of ∼3.5 μm^2^ (Figure 1e and Extended Data Figure 2). Each location was chosen to minimize overlap of the UV-A pulses. After irradiation, the cells were imaged for a further 22 hours to observe cell recovery. We recorded separate time-series images of the chlorophyll (640 nm ex./685 nm em.) and phycobilisome fluorescence (561 nm ex. /665 nm em.) every 30 minutes, along with brightfield images of the cells (Fig. 1f).

### Quantitative high-throughput image analysis

To analyze the resulting images, we used a customized computational pipeline (CyAn) that we previously developed in MATLAB (R2019a)^14^. The brightfield channel was used to identify individual cells and parameters including cell length and fluorescence intensity were measured over the duration of the movie. CyAn was also used to track individual cells and identify cell division events. Each cell was assigned a unique identifier (ID), and the ID of mother-daughter cells were recorded for each division event, creating a binary tree. A binary traversal algorithm then retrieves the lineage of cells from this tree, allowing individual cell lines to be identified over multiple generations. The type of each cell (WT or *Δcpc*) was also identified and categorized automatically. Since the *Δcpc* strain lacks phycobilisome rods, they exhibit reduced fluorescence in the phycobilisome channel compared to WT (Extended Data Fig. 3). We used this property to classify the cells based on their mean phycobilisome fluorescence intensity in the first frame.

Here we report our experimental results from three representative photodamaging conditions: low irradiation intensity (∼1.3 mW, 0.5 s exposure), high irradiation intensity (∼19.7 mW, 2 s exposure), and a condition with intermediate irradiation intensity (∼6.9 mW, 2 s exposure). The low and high irradiation conditions served as controls (Fig. 2). A qualitative assessment of the remaining 32 imaging conditions can be found in Extended Data Table 1.

**Figure 2.**
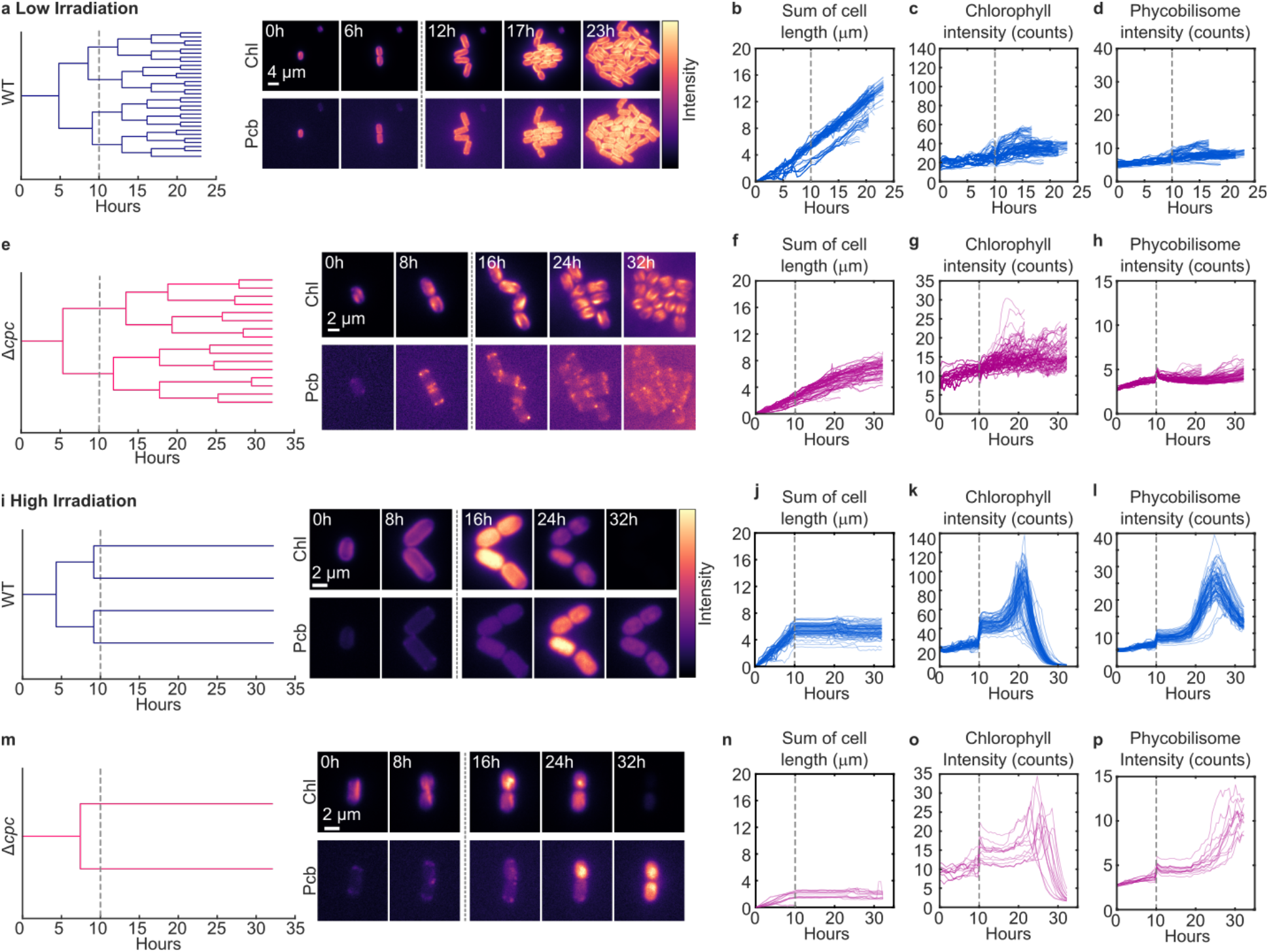
Low and high UV-A irradiance conditions. **a**, Lineage tree of a representative wild type colony in the low irradiation condition. Each branch represents a cell, and each bifurcation indicates a division event. The length of each branch is the time taken before the cell divides. Representative images of the colony at equally spaced timepoints are shown to the right of the tree. The intensity of these images is normalized to the maximum intensity of the sequence. **b**, The sum of cell lengths, **c**, the mean chlorophyll intensity (Chl), **d**, the mean phycobilisome intensity (Pcb) of all wild type lineages (n = 184) in the low irradiation movie. **e**, Lineage tree of a representative *Δcpc* colony in the low irradiation condition. **f-g**, Cell length, mean chlorophyll intensity and phycobilisome intensity of all *Δcpc* (n = 125)in the low irradiation movie. **i**, Lineage of representative wild type colony in the high irradiation condition, with corresponding data of all wild type colonies (n = 82) in the movie (**j-l**). **m**, Lineage of representative *Δcpc* colony in the high irradiation condition, with corresponding data from all *Δcpc* colonies (n = 17) in the movie (**n-p**). The dotted gray lines indicate the time when UV-A irradiation was applied.

In the low intensity condition, both strains continued to grow normally after irradiation, as shown by the representative lineage tree in Fig. 2a and in Supplementary Video 1. Note that WT has a much shorter doubling time (∼ 3.5 - 4 hours) compared to the *Δcpc* strain (6 - 7 hours). WT cells also developed into large, three-dimensional colonies at the end of the movie. Thus, tracking of these cells was stopped at the 24-hour mark because image segmentation becomes more inaccurate when applied to multi-layered colonies.

To quantify growth, the ancestors of each cell present at the end of the movie were identified by tracing the binary tree representing the single-cell lineage information back in time. We then computed the difference in cell length at each timepoint of the entire lineage. The cumulative sum of these differences was then calculated and plotted to show cell-lineage-specific growth over time (Fig. 2b). In the low irradiation condition, all cell lineages showed continuous growth. The average chlorophyll and phycobilisome fluorescence intensity of each cell in the lineages were also measured (Fig. 2c and d). There was no significant change in these intensities after irradiation in WT. We found similar results when repeating the analyses on the *Δcpc* colonies, although a small rise in phycobilisome was observed immediately following UV-A stimulation before returning to baseline levels (Fig. 2e-h).

In comparison, both WT and Δ*cpc* strains stopped growing immediately following exposure to high intensity irradiation (Fig. 2i and m, and Supplementary Video 2). This dosage of UV-A likely generated irreversible damaged that could not be repaired, as no instance of growth following the irradiation event could be identified (Fig. 2j and n). We also found that the chlorophyll intensity increased over time, reaching a maximum at ∼11 hours after irradiation for the wild type cells (Fig. 2k) and ∼15 - 19 hours after irradiation for the *Δcpc* cells (Fig. 2o). The phycobilisome intensity also increased over time, reaching a maximum at ∼15 hours after irradiation for the wild type cells (Fig. 2l) and ∼20 hours for the *Δcpc* cells.

Increased chlorophyll fluorescence intensity before a rise in phycobilisome fluorescence is consistent with our previous findings of mechanically-induced photodamage in cyanobacteria and further indicates that PSII photochemistry is disrupted prior to detachment of membrane-bound phycobilisomes and dissipation of excess energy as fluorescence under UV-A light-induced photoinhibition^18^. Notably, complete bleaching of chlorophyll fluorescence is observed by the 32 hour timepoint in WT and Δ*cpc* cells. Phycobilisome fluorescence is more stable but eventually bleaches in WT and appears to begin to bleach in some cell traces in Δ*cpc*, albeit with slower kinetics.

### Asymmetric response of cells to photodamage

Exposure of cells to intermediate levels of UV-A resulted in asymmetric survival of sister cells in WT and *Δcpc* strains (Fig. 3; Supplementary Video 3). As before, we calculated the sum of lengths for each cell present at the time of irradiation. This information was then used to classify the outcome of each colony into one of three categories: ‘growing’ - all cells continued to grow in length, ‘stopped growing’ - all cells stopped growing after irradiation, and ‘asymmetric survival’ - one cell continued to grow while its siblings stopped growing.

**Figure 3.**
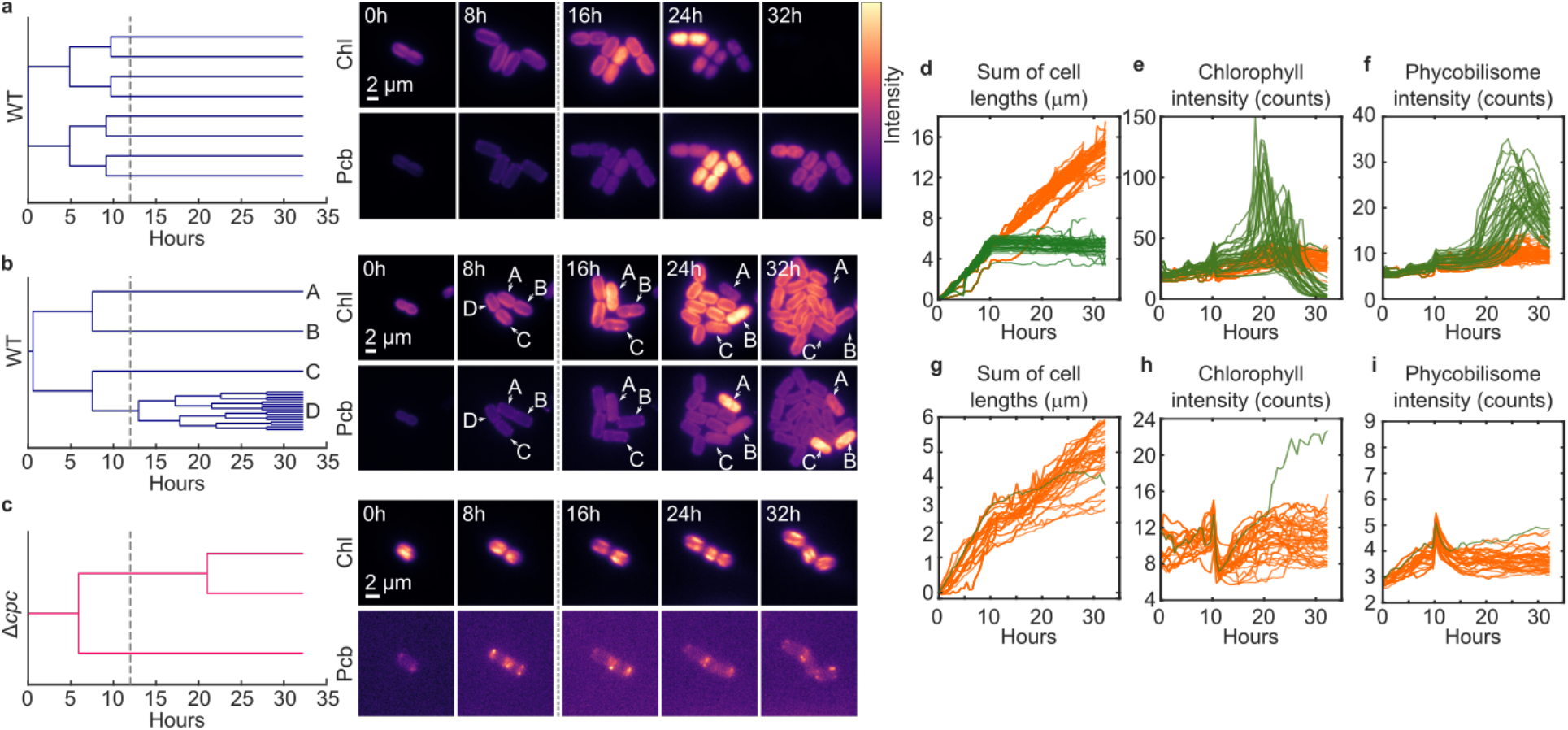
Asymmetric survival in response to intermediate UV-A irradiation. **a**, Representative lineage of a WT colony, where all cells stopped growing after irradiation. **b**, Representative lineage of a wild type colony showing asymmetric survival. **c**, Representative lineage of a *Δcpc* colony. Despite one of the branches not dividing over the duration of the time-course, the cell still appears to be growing in length. **d-f**, Sum of cell lengths, chlorophyll, and phycobilisome intensity of wild type lineages (n = 139). **g-i**, Sum of cell lengths, chlorophyll, and phycobilisome intensity of *Δcpc* lineages (n = 44). In d-i, the green traces are cells which stopped growing, while the orange traces indicate cells which continued to grow after irradiation. The dotted gray lines indicate the time when UV-A irradiation was applied.

For the WT strain, the state of each cell at the end of the movie was determined by choosing a minimum threshold for the sum of cell lengths. WT cell lineages which had a combined length of > 8.5 μm were classified as ‘growing’, while those with combined lengths less than this value were classified as ‘stopped growing’. Of the 19 WT colonies analyzed, 9 stopped growing (Fig. 3a), while 10 showed ‘asymmetric survival’ (Fig. 3b). For the *Δcpc* cells, the cell state was categorized using the mean chlorophyll intensity. Cells which had an intensity of less than 15 counts were classified as growing. In contrast to the WT colonies, all *Δcpc* colonies except one belonged to the ‘growing’ class after irradiation based on this criterion (Fig. 3c). The remaining colony showed ‘asymmetric survival’.

Figures 3d-i shows the sum of cell lengths, chlorophyll, and phycobilisome intensities of the cells. Cell lineages which were classified as stopped growing (green traces) showed an increase in chlorophyll and phycobilisome intensities, similar to the high irradiation control. Conversely, cell lineages which continued to grow (orange traces) did not show an increase in the chlorophyll or phycobilisome intensity, similar to the low irradiation control.

In ‘non-growing’ WT and *Δcpc* strains, we also identified the emergence of a brightly fluorescent puncta in the chlorophyll channel after irradiation (Fig. 4; Supplementary Video 3). In WT cells, the structure also exhibits phycobilisome fluorescence, which was not apparent in the Δ*cpc* mutant. We hypothesize that these puncta are associated with the PSII repair cycle as intracellular vesicles have been previously observed when imaging cyanobacteria, albeit not necessarily in connection to photoinhibitory conditions^24–26^.

**Figure 4.**
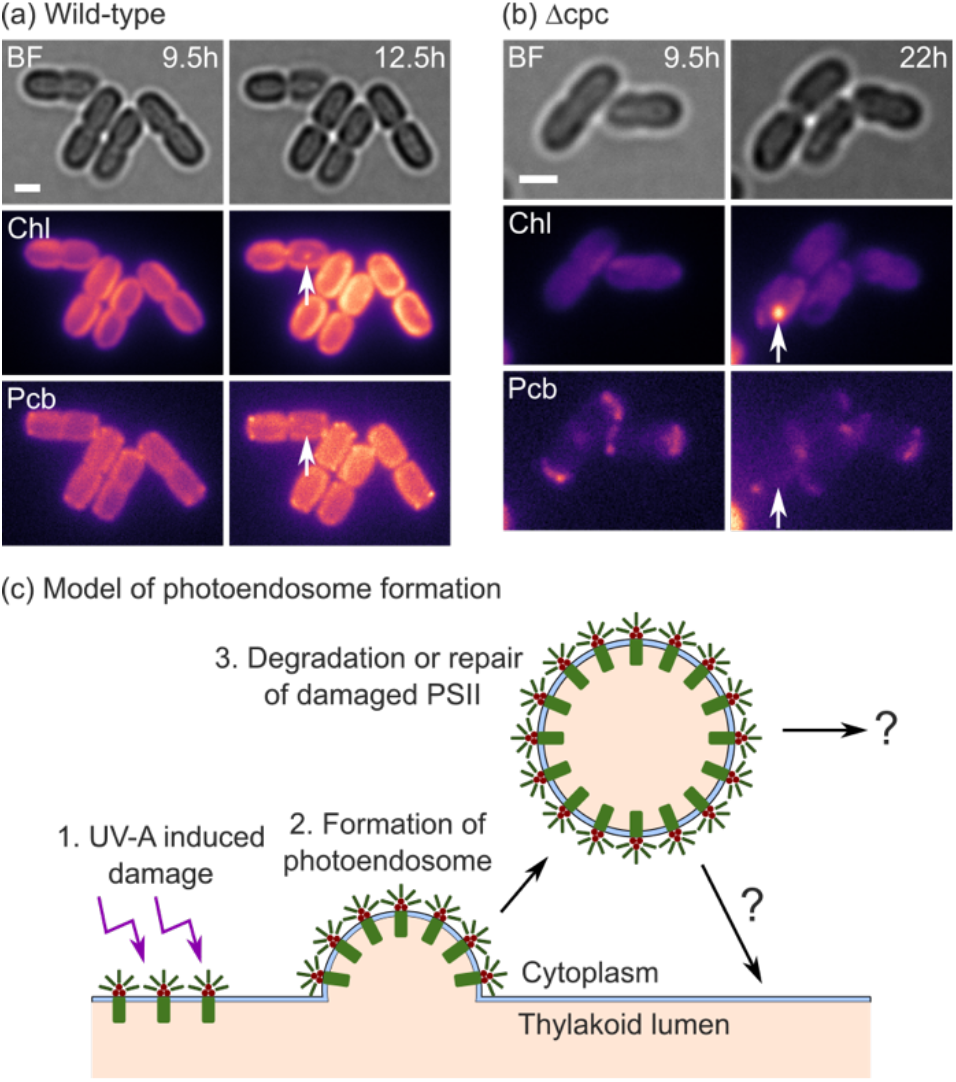
Identification and model of photoendosome formation. **a**, Representative images showing internalized vesicle-like structures (photoendosome) with chlorophyll (Chl) and phycobilisome (Pcb) fluorescence in WT cells. **b**, A similar structure was found in the only *Δcpc* cell that stopped growing after irradiation. The white arrows indicate the position of the fluorescent puncta. Scale bars are 2 μm. **c**, Proposed model of photoendosome formation after UV-A induced photodamage.

## Discussion

Here, we have demonstrated an automated imaging approach which combines long-term time-lapse microscopy and computerized image analysis to enable photoinhibition and recovery to be studied in single cell lineages. This approach has enabled observations which show that cyanobacteria growing in colonies can exhibit asymmetric survival in photoinhibitory conditions. The discovery of this asymmetric survival in sibling cells derived from a common lineage suggests that non-genetic cell-cell heterogeneity could be an underlying mechanism to survive unfavorable conditions. Non-genetic phenotypic differences, including bet-hedging, have been previously observed in other microbes, including other species of cyanobacteria^27^. An alternative explanation for asymmetric survival could also be due to genetic differences. Cyanobacteria are polyploid and other polyploid bacterial species, like the radiation resistant *Deinocooccus radiodurans* are known to pass on non-equivalent genomes to daughter cells^28–30^.

Additionally, we have observed the formation of brightly fluorescent spots in the photodamaged cells, which we hypothesize are related to PSII repair and could be mediated by the vesicle-inducing protein Vipp1. This hypothesis is based on several pieces of evidence; Recent structural studies have revealed that Vipp1, found in cyanobacteria and plant chloroplasts, and the endosomal sorting complexes required for transport-III (ESCRT-III) are members of the same superfamily of membrane remodeling proteins^31–33^. The ESCRT-III complex is known to be involved in membrane remodeling, including membrane budding from the cytoplasm. Additionally, in PCC 7002, Δ*vipp1* mutants are unable to grow photoautrophically, but can be grown photoheterotrophically in low light with the addition of glycerol as a carbon source^34^. Furthermore, Vipp1 has been shown to re-localize to foci on the thylakoid membrane and co-precipitate with stress response proteins as well as phycobilisome rod and core subunits in cyanobacteria in response to high-light exposure^35^, suggesting a role in the photodamage and repair pathway.

We propose the term “photoendosome” as a new name for this intracellular vesicle in photosynthetic organisms and organelles and present a model for its formation in response to UV-A induced photodamage (Fig. 4c). Physical separation and internalization of damaged-but-photoactive complexes, which could act as photosensitizers, may be a mechanism to prevent the catalytic spread of photodamage to adjacent healthy membranes by short-lived reactive oxygen species and other free radicals during the repair process. Further studies to determine the mechanism of photoendosome formation and asymmetric survival pattern are ongoing and may provide insight into processes critical for oxygenic photosynthesis.

## Methods

### Imaging

Imaging was carried out using a commercial mechanized microscope (Nikon TiE). Automated control of the microscope was performed by the supplied software (Nikon NIS-Elements AR, version 5.11.00, 64-bits). The microscope has two illumination sources: Excitation light was provided by a solid-state light source (Lumencor, SpectraX), while growth light was provided by a light engine (Lumencor, LIDA). The sample stage of the microscope was enclosed by a cage incubator (Okolab) which provides temperature control. For these experiments, the samples were grown in air at a temperature of 37 °C. To ensure thermal stability, the temperature unit was turned on for at least an hour before imaging commenced. Furthermore, after the imaging dish was placed on the translational stage of the microscope, it was left for 30 minutes to allow the samples to reach thermal equilibrium before the microscope was focused. This process was important in maintaining focus throughout the experiment. During this initial incubation period, the cells were illuminated with red light at an intensity of ∼157 μmol photons m^2^ s^-1^. For the proof-of-principle experiment, 35 different locations on the same agar pad were pre-selected. The microscope was then programmed to image each location using the Jobs module in the Nikon NIS-Elements software. During the irradiation step, the microscope was programmed to illuminate the cells at each location with a pulse of UV-A light with different durations and intensities. The exposure time was varied between 0.5 - 3.5s in steps of 0.5 s, while the intensity setting was varied between 5 - 85% in steps of 20%. All images were taken using a 100x oil immersion objective (Nikon, CF160 Plan Apochromat, NA = 1.45). At each timepoint, a set of three images were acquired. The chlorophyll fluorescence was imaged using the Cy5 filter set (peak excitation 645 nm, peak emission 705 nm, 2% laser power of 0.368 mW). The phycobilisome fluorescence was imaged using the RFP filter set (peak excitation 555 nm, peak emission 665 nm, 40% laser power of 18.2 mW). Brightfield image of the cells were also acquired for image segmentation and to validate our observations.

To measure the UV-A light power, we closed the internal aperture of the microscope, so the illuminating beam was approximately the same area as the field of view (∼0.13 mm by 0.13 mm). We then used a high intensity power meter (Argolight, Argo-POWER) to measure the illumination power vs laser setting of the NIS-Elements software (Extended Data Figure 2).

### Image analysis

The WT and *Δcpc* cells were identified using the phycobilisome fluorescence intensity of the cells in the first frame of the movie. We set a threshold intensity of 400 counts; Cells with phycobilisome fluorescence larger than this were classified as WT, while the rest were classified as *Δcpc*. This threshold was chosen by manually identifying several cells from each movie, then plotting the histogram of the mean phycobilisome fluorescence, as shown in Extended Data Figure 3.

As mentioned above, we used a customized pipeline CyAn to track individual cell lineages throughout the time-lapse movies. Using the cell IDs from the tracking code, the lineage of each cell could be traced. The algorithm worked as follows. First, cells which did not have daughters (also called leaf nodes) at the end of the movie were identified. For each identified cell, we traversed backwards through the mother-daughter tree to determine its lineage (Extended Data Figure 4).

For the traces shown in Figures 2 and 3, we cleaned the data to remove traces containing tracking or segmentation errors. In particular, traces from cell lineages which were not tracked through to the end of the movie were removed. We also removed plots with large variations in cell lengths or intensity as these are indicative of segmentation or tracking errors, as well as traces from cells which grew out of the plane of focus. Code used to sanitize the data is provided in the Github repository listed below.

## Data availability

The datasets generated and analyzed during the current study are available from the corresponding author on reasonable request.

## Code availability

Code is available at https://github.com/jwtay1/photoinhibition-cyanobacteria

## Reporting summary

Further information on research design is available in the Nature Research Reporting Summary linked to this paper.

## Acknowledgements

This study was financially supported in part by the U.S. Department of Energy (DOE) DE-SC0019306 and DE-SC0020361to J.C.C.

## Author contributions

J.C.C. conceived the project and obtained funding. J.W.T. and J.C.C. designed the experiments. J.W.T. carried out the optical microscopy, developed the analysis code, and performed data analysis with input from J.C.C.. J.W.T. and J.C.C. wrote the manuscript.

## Extended Data

**Extended Data Figure 1.**
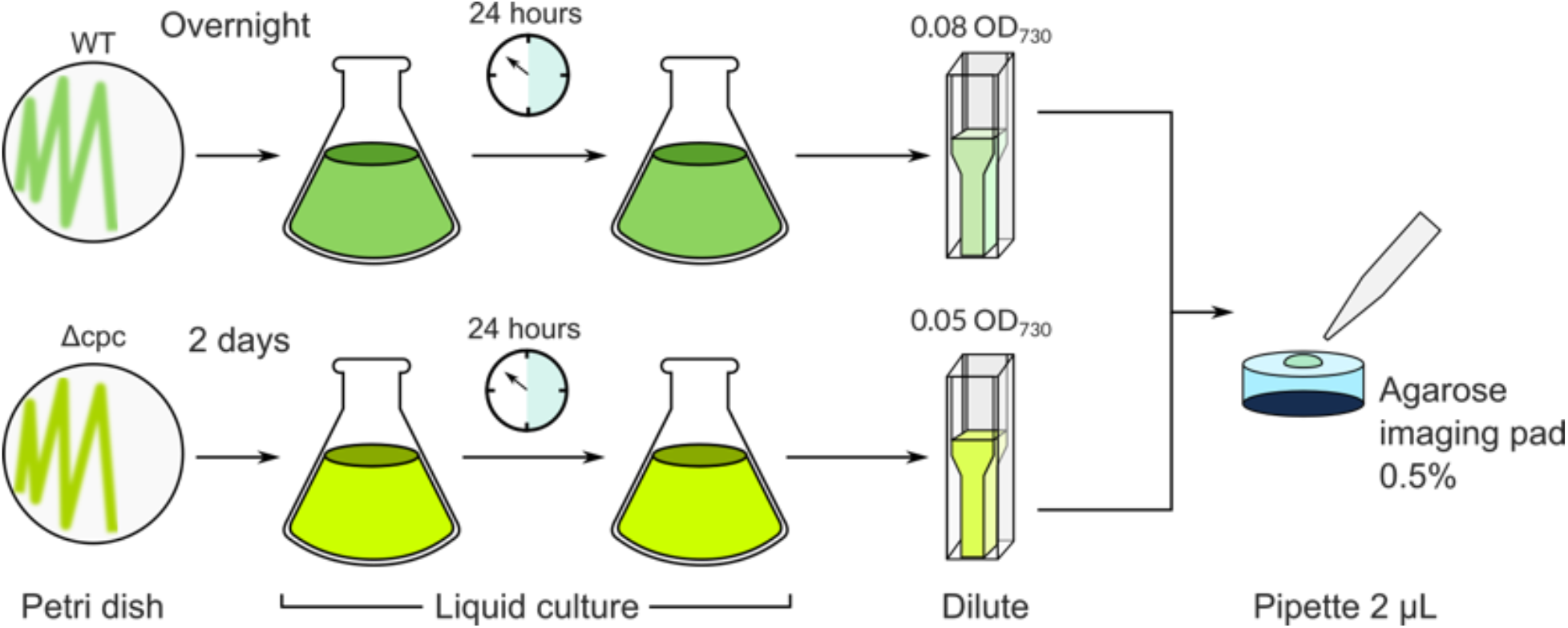
Preparation of WT and Δ*cpc* strains for co-imaging. WT grows faster than the Δ*cpc* strains. Thus, pre-cultures were started from solid media and acclimated for 24-48 h prior to dilution into experimental cultures grown for 24 h. Cells were diluted to the same cell density prior to mixing 1:1 and transfer to 0.5% agarose imaging pad prepared with standard A+ media. The optical densities of the cultures were selected to account for slight differences in the OD730nm/cell number between strains, likely arising from different cell sizes and subcellular compositions.

**Extended Data Figure 2.**
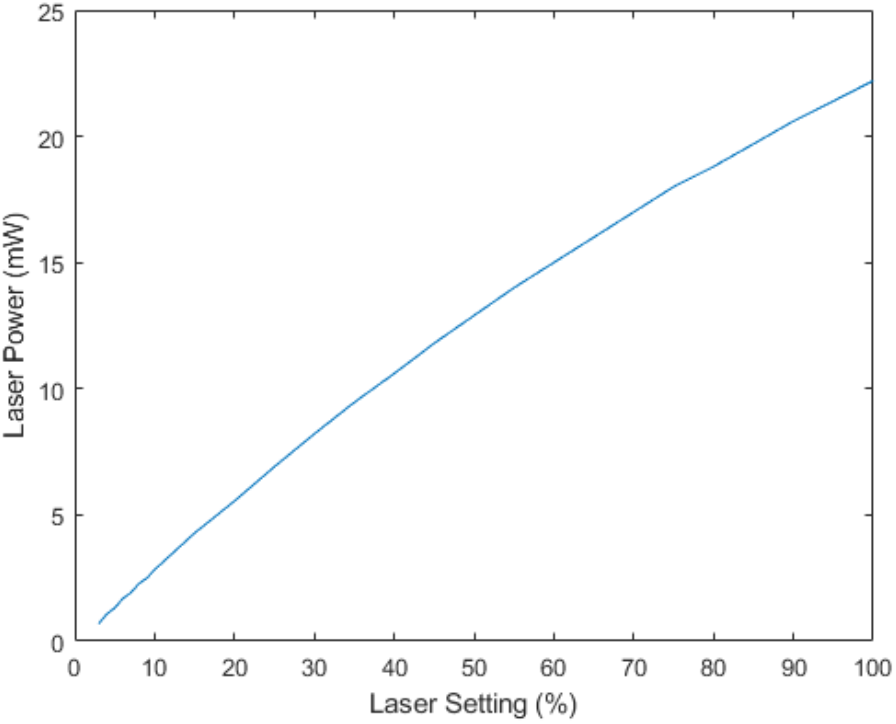
Measured UV-A power. The power of the UV-A light source was measured by closing the microscope aperture to match the field of view. The total illumination intensity was then measured using a power meter (Argolight, Argo-POWER).

**Extended Data Figure 3.**
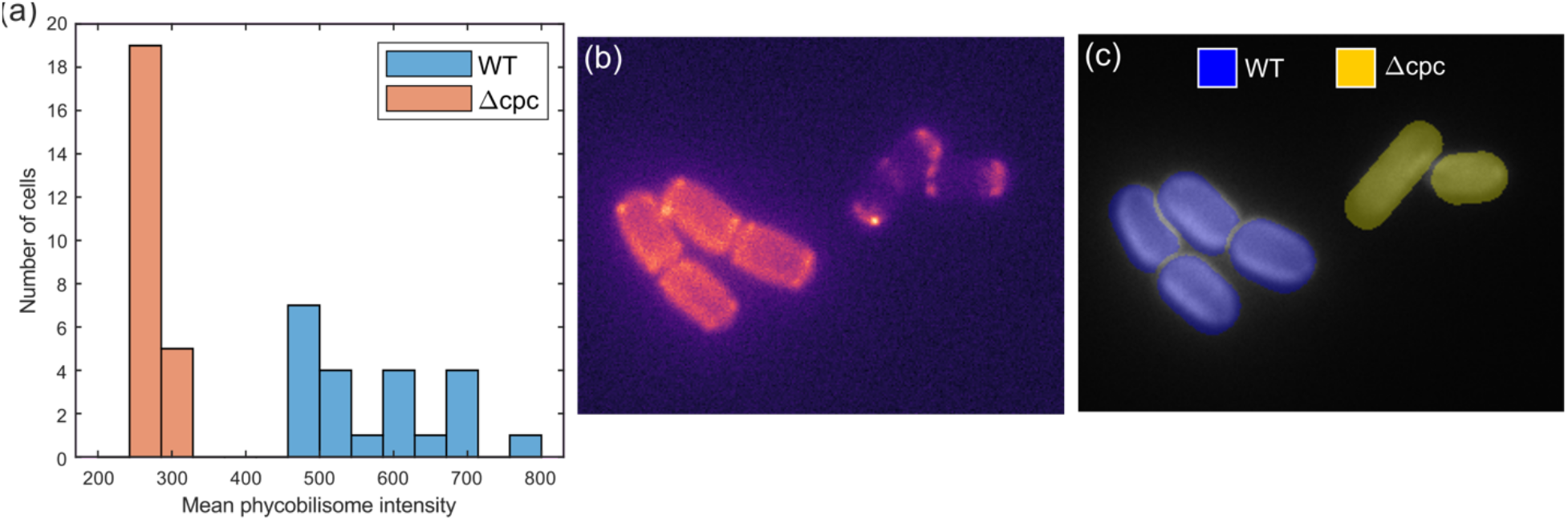
Choosing a threshold for cell identification. **a**, We manually identified cells of each type from the three movies (total WT = 24, total *Δcpc* = 22), then plotted a histogram of the mean phycobilisome intensity of each cell. Using this data, we chose a threshold intensity of 400 counts to classify the rest of the cells. **b**, Representative image showing the phycobilisome fluorescence. **c**, Cell labels overlaid over the chlorophyll channel. WT cells are colored blue, while the *Δcpc* cells are colored yellow.

**Extended Data Figure 4.**
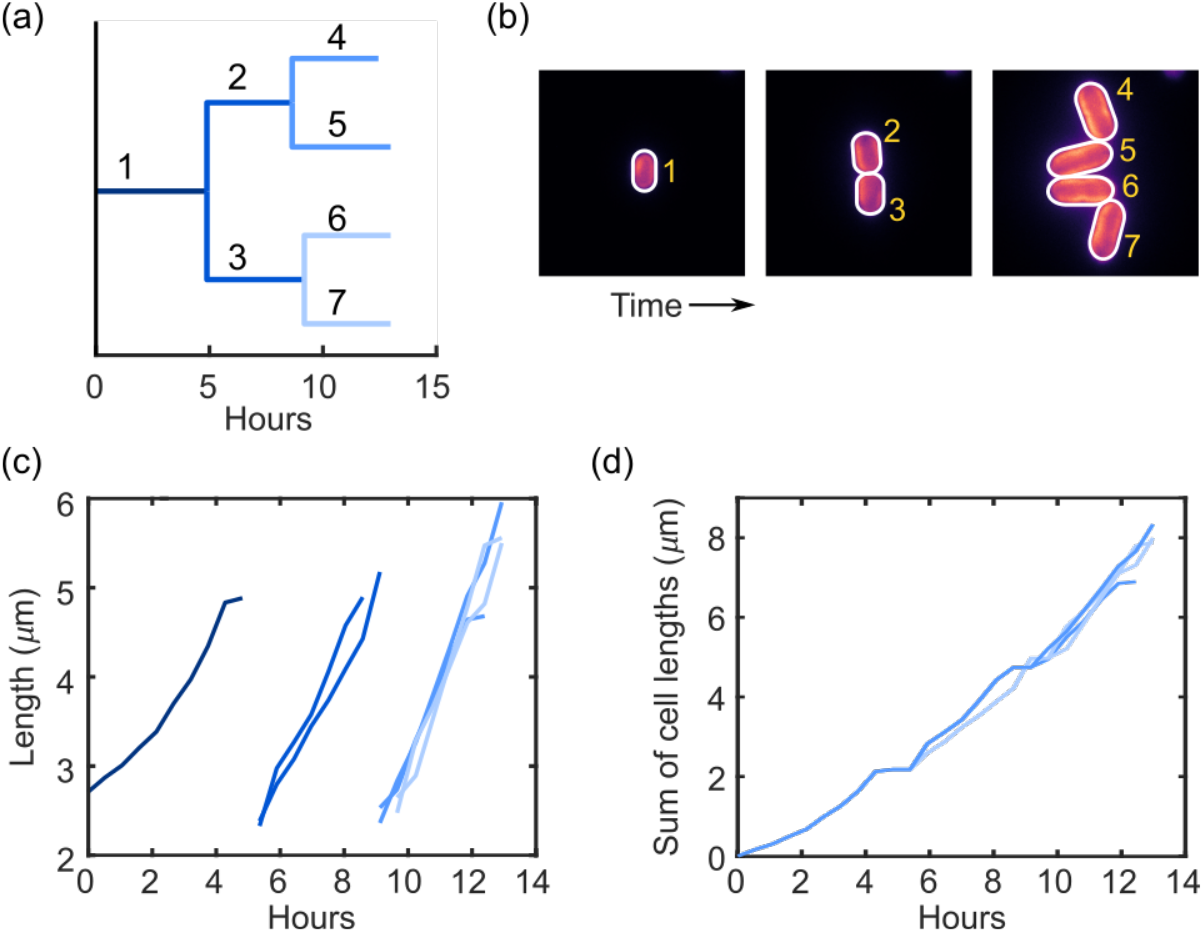
Lineage tracing. **a**, Lineage tree of a representative colony. **b**, Individual frames captured from the movie of the colony. **c**, The length over time of each cell in the movie. **d**, The resulting sum of cell lengths.

**Extended Data Table 1.**
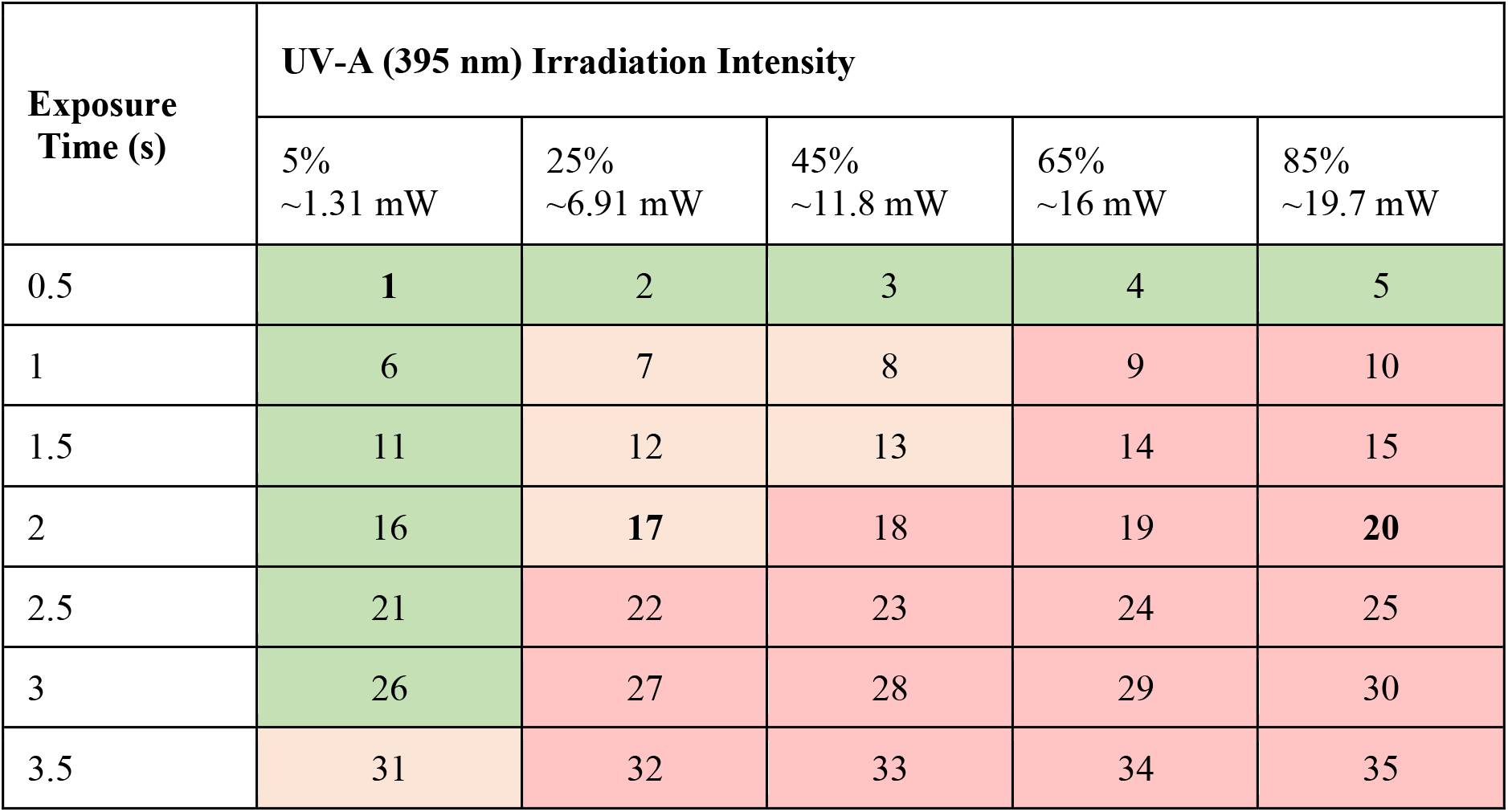
Qualitative assessment of colony outcomes. We manually evaluated all 35 irradiation conditions and classified the outcome as all cells grew (green boxes), all cells stopped growing (red boxes), and asymmetric survival (orange boxes). Results from conditions 1 (low irradiation), 20 (high irradiation), and 17 (intermediate irradiation) were used in this article.

## Supplementary Videos

**Supplementary Video 1** | **Low irradiation intensity**. Brightfield (left), chlorophyll fluorescence (middle), and phycobilisome fluorescence (right) channels, showing a representative WT colony (top) and *Δcpc* colony (bottom) exposed to low light irradiation. The color scale for each channel is normalized the maximum intensity of the movie.

**Supplementary Video 2** | **High irradiation intensity**. Brightfield (left), chlorophyll fluorescence (middle), and phycobilisome fluorescence (right) channels, showing a representative WT and *Δcpc* colonies exposed to high light irradiation. Note that all cells stop growing after irradiation (frame 19), as well as exhibiting increased chlorophyll and phycobilisome fluorescence. The color scale for each channel is normalized the maximum intensity of the movie.

**Supplementary Video 3** | **Intermediate irradiation intensity**. Brightfield (left), chlorophyll fluorescence (middle), and phycobilisome fluorescence (right) channels, showing a representative WT colony (top) and *Δcpc* colony (bottom) exposed to intermediate irradiation. Note that colonies can be seen here which show both arrested growth, continued growth, and asymmetric survival. A *Δcpc* cell in the middle of the frame also shows the formation of a bright puncta, here termed a ‘photoendosome’. The color scale for each channel is normalized the maximum intensity of the movie.

**Supplementary Video 4** | **Tracking individual cells**. Representative video showing results of segmentation and tracking on the intermediate irradiation intensity movie. Cells are identified by green outlines. Unique IDs issued by the tracking code is shown by the yellow numbers. The ID remains constant until the cell divides and the daughter cells are then assigned new IDs.

